# The cross-domain functional organisation of posterior lateral temporal cortex: Insights from ALE meta-analyses of seven cognitive domains spanning 9515 participants

**DOI:** 10.1101/2022.01.18.476749

**Authors:** Victoria J. Hodgson, Matthew A. Lambon Ralph, Rebecca L. Jackson

## Abstract

The posterior lateral temporal cortex is implicated in many verbal, nonverbal and social cognitive domains and processes. Yet without directly comparing these disparate domains, the region’s organisation remains unclear; do distinct processes engage discrete subregions, or could different domains engage shared neural correlates and processes? Here, using activation likelihood estimation meta-analyses, the bilateral posterior lateral temporal cortex subregions engaged in seven domains were directly compared. These domains comprised semantics, semantic control, phonology, biological motion, face processing, theory of mind, and representation of tools. Whilst phonology and biological motion were predominantly associated with distinct regions, other domains implicated overlapping areas, perhaps due to shared underlying processes. Theory of mind recruited regions implicated in semantic representation in the left hemisphere, tools engaged semantic control areas, and faces engaged subregions for biological motion and theory of mind. This cross-domain approach provides insight into how posterior lateral temporal cortex is organised and why.

## INTRODUCTION

The posterior lateral temporal cortex (pLTC) is a “Bermuda triangle” of cognitive function. The region, or subregions within, are implicated in a wide array of disparate domains, from aspects of language processing through to understanding intentional action, raising questions about its organisation. Is this brain area composed of many tessellated, discrete subregions each subserving different functional domains, or is pLTC responsible for a smaller number of core cognitive processes that underpin a range of domains? Focusing within a single cognitive domain, researchers may miss the clues as to a particular region’s function that could be provided by the region’s involvement in other domains. Here we take a broader view. In the present study, a series of activation likelihood estimation (ALE) meta-analyses were used to compare and contrast activation systematically across seven different cognitive domains commonly associated with bilateral pLTC. This large, cross-domain meta-analysis investigated how multiple different cognitive processes are supported by pLTC and determined the principles underlying the functional organisation of this area.

The functions ascribed to the pLTC (here defined as including posterior lateral regions, dorsally up to the temporo-parietal junction, TPJ, and excluding the basal surface of the temporal lobe) are numerous and diverse. The pLTC has been implicated extensively in semantic cognition, the representation and flexible use of our multimodal conceptual knowledge ^1,2^. Following the recognition that posterior brain damage can result in a semantic control deficit ^3–6^, meta-analyses have highlighted a particular role for pMTG in semantic control ^7,8^. However, pLTC regions are also implicated in semantic representation, both as a whole and in the representation of specific semantic categories, such as tools (defined as manipulable man-made objects) ^9–13^, actions more generally ^14,15^, faces ^16–18^ and bodies or body parts. Indeed, a portion of the pMTG known as the extrastriate body area is proposed to be specialised for the visual detection of bodies and body parts ^19–22^, and pSTS is implicated in the low level detection of biological motion across a broad range of real and point-light display stimuli ^23,24^, including bodies, faces, expressions, and mouth and eye movements ^18,24–31^. Beyond semantics, the pLTC is implicated in phonological processing, particularly in STG and the ventral aspects of inferior parietal cortex which are included in the scope of the current investigation ^32–35^. Indeed, damage around the posterior Sylvian fissure can cause conduction aphasia, characterised by impaired repetition, naming difficulties and phonemic paraphasias, despite good comprehension ^36–39^. Regions within the pLTC are also considered important for social cognition, an umbrella term for a collection of processes such as empathy, interpreting intentional actions, and false belief understanding ^23,40,41^. In particular, the TPJ is considered a crucial region for understanding the mental states of others, or ‘theory of mind’, including classic false belief tasks ^40,42,43^.

To date, the regions implicated in these myriad domains have not been systematically compared to one another. As such, it is not clear to what extent overlapping pLTC regions are implicated in different domains; for instance, is the same region of pMTG critical for semantic control, the representation of tools, and the detection of body parts? In part, this is due to the imprecision in anatomical labelling, making it difficult to know whether researchers in disparate areas of cognitive neuroscience are referring to the same, overlapping, or entirely distinct regions when using the same terminology, or indeed, if they are referring to the same regions when using different terminology. The TPJ is a prime example of this; the label ‘temporo-parietal junction’ is ill-defined and does not map precisely to anatomical features ^44^. As such it is variously used to refer to parts of the pSTG, angular gyrus (AG), and supramarginal gyrus ^11,45^. It is also likely that these established anatomical labels are too coarse to capture subregions within the posterior temporal cortex. Furthermore, functional involvement and activation patterns do not always respect neat anatomical boundaries.

To combat these difficulties, the present study used a series of ALE meta-analyses, within a pLTC ROI, as a tool for direct, statistical comparison across the key domains associated with this region, including semantics, semantic control, phonology, representation of tools, representation of faces, perception of biological motion and theory of mind. This allowed us to elucidate the functional organisation of pLTC by investigating whether different functions recruit the same areas – and as such may rely on common underlying neural processes and computations, about which we can begin to form some hypotheses – or whether they engage distinct, neighbouring subregions.

## METHODS

### Definition of ROI

To cover the pLTC, including the ill-defined TPJ, an ROI was constructed consisting of the posterior portion of ITG, MTG and STG along with the most inferior portions of parietal cortex. As studies and domains highlighting the role of the ‘TPJ’ may or may not be referring to the posterior lateral temporal cortex proper, a full TPJ region was included, allowing identification of the region in question, and comparison to other domains, irrespective of whether the region identified is strictly temporal or parietal. The basal temporal lobe was excluded as its complex organisation has already received extensive discussion and is not the focus of the present investigation ^27,46–52^. The ROI was constructed using the automated anatomical labelling (AAL) anatomical atlas ^53^ taken from MRIcron (http://www.nitrc.org.projects/mricron) in MNI space. The ROI was bounded dorsally by a horizontal plane oriented to the intraparietal sulcus on the lateral surface (z=42 in MNI space), anteriorly by a line beginning at Heschl’s gyrus on the lateral surface (y=-19, z=6 in MNI space) and progressing approximately perpendicular to the Sylvian fissure, ventrally by the boundary between the inferior temporal gyri and the fusiform gyrus, and posteriorly by the boundary with the occipital lobe as defined by the AAL atlas. The left hemisphere view of the ROI is displayed in Figure 1 and the right hemisphere view in Figure 3.

**Figure 1.**
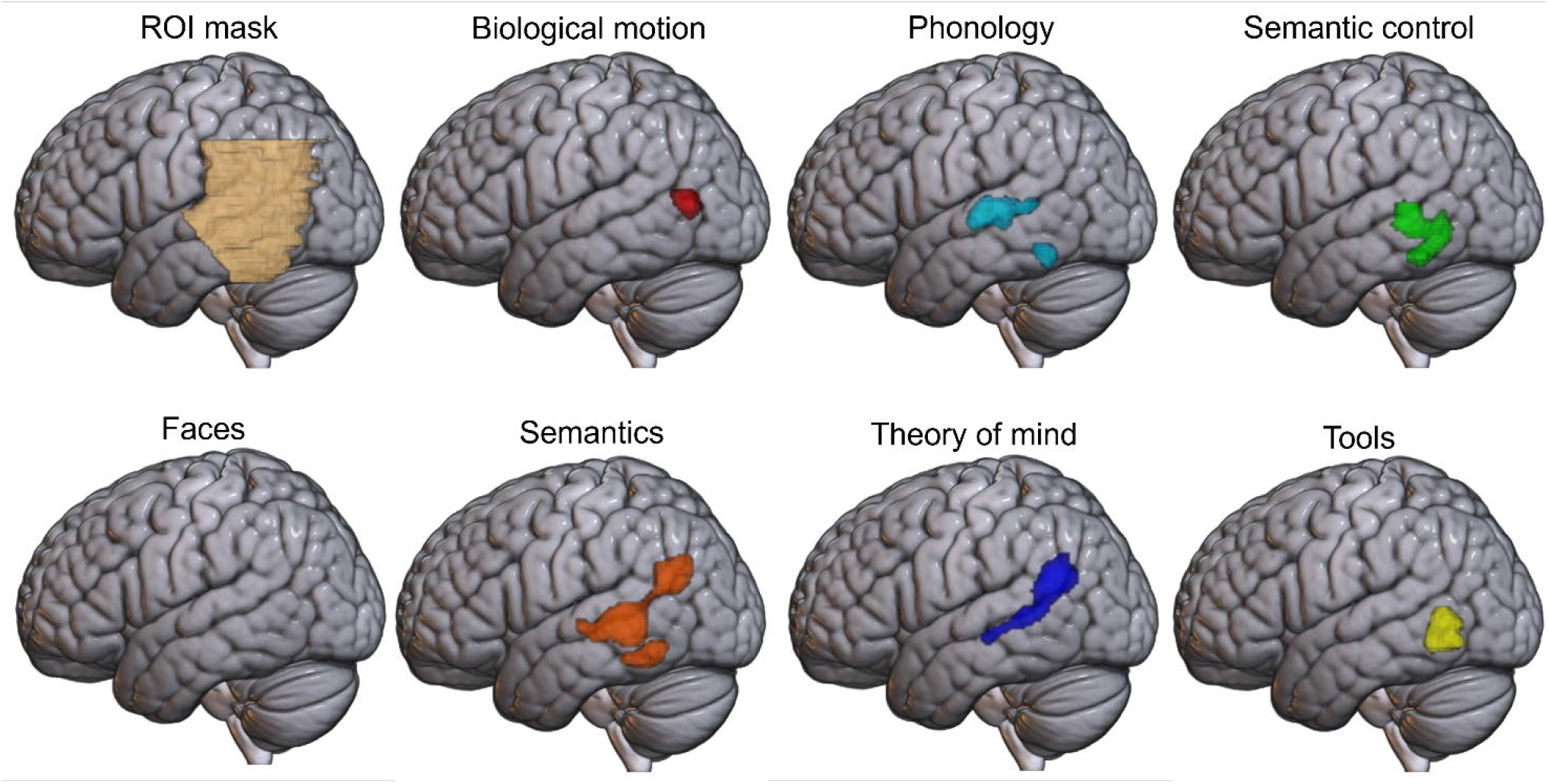
Activation likelihood estimation maps for each of the seven domains, showing clusters of consistent activation across studies in the left hemisphere, at a voxel-level threshold of p<.001. Note, no voxels were identified in the face domain. Top left: the region of interest mask used for all analyses, shown in the left hemisphere only.

### Meta-analyses

Independent ALE analyses, restricted to the ROI, were completed for each of the seven domains, before comparison between domains. Studies for each domain were sourced from one or more existing meta-analyses, in order to ensure that accepted, peer-reviewed operationalisations of each domain were used and directly compared to each other for the first time, and to maximise the breadth of the domains that were included. For each study matching the inclusion criteria, each peak was assessed for overlap with the ROI and only peaks within the ROI were included.

#### Inclusion and exclusion criteria

Analyses included only peer-reviewed English language articles, describing task-based fMRI and PET studies that reported peak coordinates of a univariate contrast in standard (MNI or Talairach) space across the whole brain and focused on a young healthy adult sample (below 40 years old). Contrasts were excluded if they focused on patients, clinical trials or individual differences (e.g., age, gender, native language). Within each included article, wherever multiple task contrasts were reported for the same participant sample, all the peak activation coordinates were analysed as a single contrast, following the recommendation from Mueller et al. ^54^. Where domains could include overlapping content that may explain recruitment of the same region (for instance, the presentation of faces in studies assessing theory of mind), relevant studies were either excluded *a priori* or, if this resulted in a large reduction in sample size, the analysis was performed with and without their removal to maximise power whilst aiding the interpretation of the between-domain comparison.

##### Semantics

To identify the pLTC regions recruited for semantic cognition across categories and processes, a general semantic contrast was included. Studies were sourced from a recent meta-analysis by Jackson ^8^, which included 272 verbal and nonverbal contrasts published between 1992 and 2019 that specifically compared a semantic condition with a non-semantic (or less semantic) condition. These studies could include individual semantic categories, such as faces, compared to a baseline but did not contrast different categories of concepts. All contrasts involving tools were removed to allow uncontaminated comparison with the tool domain. This resulted in the inclusion of 580 foci across 204 experiments after restriction to the pLTC ROI.

##### Semantic control

A further 126 contrasts assessing semantic control, in particular, were sourced from Jackson ^8^. These contrasted high over low semantic demands, using a range of manipulations, including association strength, competitor interference, and homonym ambiguity. These comparisons recruit a subset of the regions responsible for semantic cognition (dissecting semantics into areas responsible for semantic control and semantic representation). All contrasts involving tools were removed. Within the ROI, there were 104 foci across 41 experiments.

##### Phonology

Studies were sourced from a meta-analysis by Hodgson et al. ^55^, which identified studies from the Vigneau et al. ^34^ and Humphreys & Lambon Ralph ^56^ meta-analyses and performed a literature search extending the timespan of the studies included to April 2021. Tasks included both passive listening and active judgements. The studies contrasted either phonological with semantic or orthographic judgements, or phonological with non-phonological stimuli (e.g., visual perception tasks). Contrasts were excluded if the phonological task contained words or other overtly semantic stimuli. From an initial pool of 82 papers published between 1992 and 2021, 207 foci across 64 experiments were included.

##### Theory of mind

Studies were sourced from a meta-analysis by Molenberghs et al. ^57^, resulting in a pool of 132 papers published between 1999 and 2014. This meta-analysis included both affective and cognitive theory of mind tasks, implicit and explicit task instructions, and varying stimuli (photographs, cartoons, stories, games, videos, animations). Commonly used tasks included the Reading the Mind in the Eyes task for mental state evaluation ^58^, the false belief over false photograph task ^59^ or similar belief over physical reasoning, and social over non-social games or animations. For the full dataset, 329 foci across 113 experiments were included. To aid precise interpretation of the regions initially implicated in both theory of mind and face processing, a subset of the dataset was analysed further, in which contrasts including face stimuli were removed. This reduced dataset included 286 foci across 90 experiments.

##### Biological motion

Studies were sourced from a meta-analysis by Grosbras et al. ^60^, which included 110 papers published between 1996 and 2010. The small number of contrasts that did not include full motion, but only static or implied motion, were removed. All contrasts compared biological motion over scrambled motion, non-biological motion, or static images. The biological motion stimuli included point light displays and moving body regions such as the hands or face, and most studies involved passive viewing. Any contrasts that included tool manipulation were removed, to eliminate overlap with the tool domain. Trials including manipulation of non-tool items (e.g., grasping a ball or similar object) were included. The full dataset comprised 223 foci across 53 experiments. In addition, a subset of this dataset was analysed, in which any contrasts with stimuli that included faces – real or animated – were removed; this subset comprised 151 foci across 40 experiments.

##### Faces

Studies were sourced from two meta-analyses by Müller et al. ^61^ and Eickhoff et al. ^62^, yielding a total of 139 papers published between 1992 and 2015. Contrasts included faces over non-face objects (e.g., houses). Therefore, while stimuli could include both emotional and neutral faces, none explicitly assessed the effect of emotion by contrasting emotional over neutral faces. The full dataset included 85 foci from 41 experiments within the pLTC. A subset of this dataset with all contrasts featuring emotion evaluation tasks removed, was analysed, with a total of 48 foci from 24 experiments, with most remaining studies employing gender evaluation or identity judgement tasks.

##### Tools

Contrasts for the tools domain were drawn from three, partially overlapping meta-analyses by Ishibashi et al. ^63^, Humphreys & Lambon Ralph ^56^ and Chen et al. ^64^, resulting in a total pool of 76 papers published between 1996 and 2013. All contrasts included the presentation of tool or tool-related stimuli over the presentation of non-tool stimuli, including other semantic categories, such as animals or faces, and non-semantic items such as scrambled images. Contrasts typically included photographs of tools over non-tools, but in some cases focused on tool sounds, action verbs and motor imagery specifically related to tools. The small number of experiments with dynamic stimuli were removed to reduce conflation with the biological motion domain. The final dataset included 116 foci from 41 experiments.

#### Activation likelihood estimation

Meta-analyses were performed using activation likelihood estimation (ALE) in GingerALE version 3.0.2 using the command line (https://brainmap.org/ale/; ^62,65–67^; code included in Supplemental Materials 1). ALE is a meta-analytic technique that maps the statistically significant convergence of activation probabilities between experiments considered to reflect similar processes. This is achieved by modelling all foci for each experiment as Gaussian probability distributions, with the full width at half maximum (FWHM) of each Gaussian being determined by the sample size of the study (i.e., larger samples result in less uncertainty of the peak’s location and a narrower distribution). This results in a modelled activation map for each experiment included in the analysis. The union of these maps is then calculated to produce an ALE score in each voxel, with each ALE score representing the probability of activation being present at that given voxel in a study.

All analyses were performed in MNI space and restricted to the ROI. Only foci that fell within the ROI were included in each contrast. The ROI was used as a mask for all statistical analyses, restricting the possible locations where a peak would be expected to fall by chance, in line with Müller et al. ^54^. This allows assessment of whether the peak coordinates generated across studies are more consistently clustered than would be expected by chance, within the volume of the ROI. ALE scores were thresholded with a voxel-wise p-value of .001. Cluster-level family-wise error correction at a p-value of .001, with 10000 permutations, was then applied to determine the minimum significant cluster size and remove non-significant clusters.

In addition to the main analyses for each domain, pairwise contrast analyses were performed, with conjunction and subtraction analyses revealing the distinct and shared areas across each pair of domains. Contrasts are only presented in the main text for pairs of domains with overlapping activations, with the remainder presented in the Supplementary Materials (Supplementary Figures 1-13). These contrast analyses similarly used the ROI mask, a p-value of .001 with 10000 permutations and a minimum cluster volume of 20mm^3^.

## RESULTS

Activation peaks for each of the seven domains are provided in Table 1. As laterality appears an important organisational factor, the results are presented separately for each hemisphere followed by a consideration of the effect of laterality.

**Table 1.**
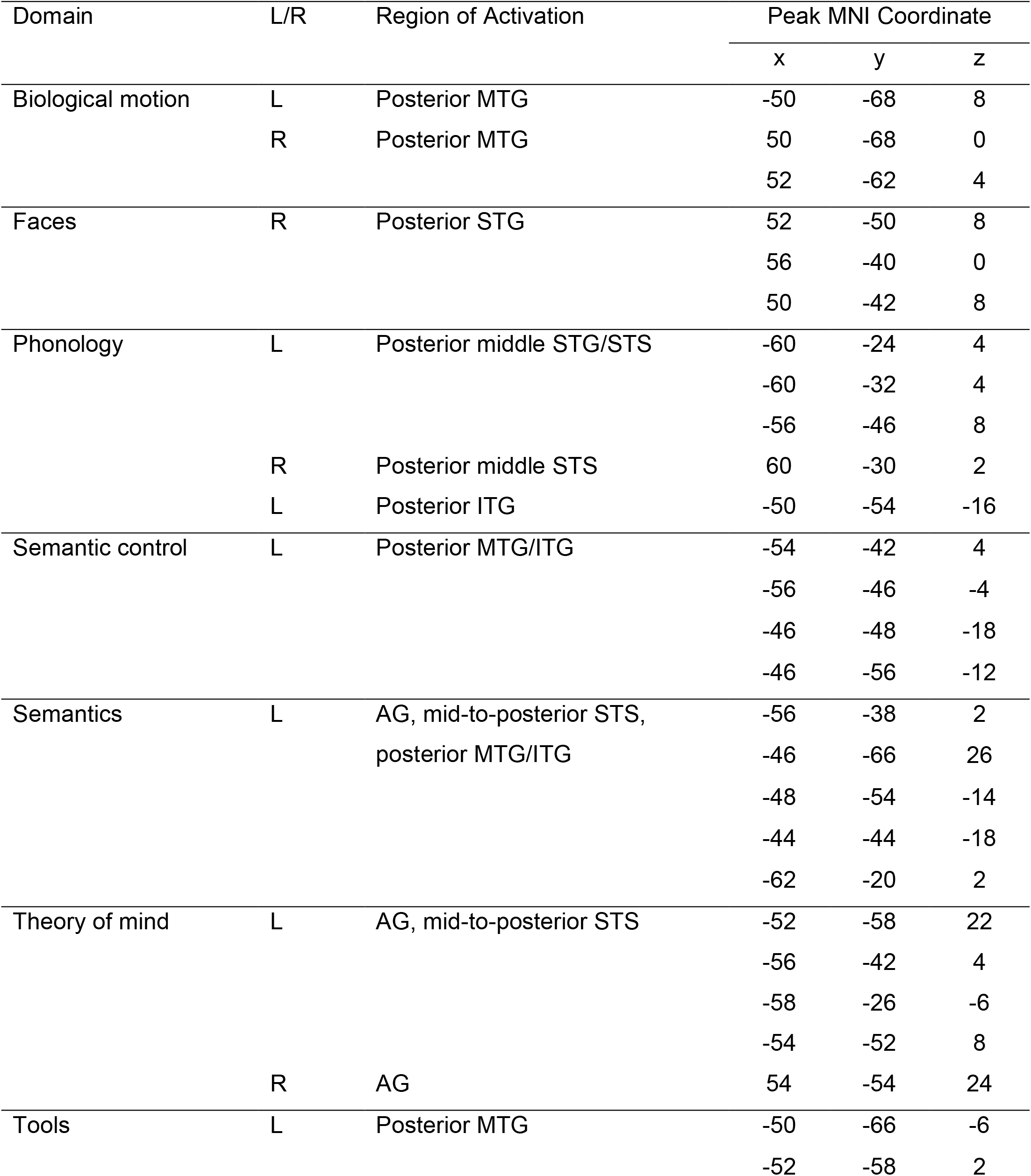
Activation likelihood estimation across all domains

| Domain | L/R | Region of Activation | Peak MNI Coordinate |  |  |
| --- | --- | --- | --- | --- | --- |
|  |  |  | x | y | z |
| Biological motion | L | Posterior MTG | -50 | -68 | 8 |
|  | R | Posterior MTG | 50 | -68 | 0 |
|  |  |  | 52 | -62 | 4 |
| Faces | R | Posterior STG | 52 | -50 | 8 |
|  |  |  | 56 | -40 | 0 |
|  |  |  | 50 | -42 | 8 |
| Phonology | L | Posterior middle STG/STS | -60 | -24 | 4 |
|  |  |  | -60 | -32 | 4 |
|  |  |  | -56 | -46 | 8 |
|  | R | Posterior middle STS | 60 | -30 | 2 |
|  | L | Posterior ITG | -50 | -54 | -16 |
| Semantic control | L | Posterior MTG/ITG | -54 | -42 | 4 |
|  |  |  | -56 | -46 | -4 |
|  |  |  | -46 | -48 | -18 |
|  |  |  | -46 | -56 | -12 |
| Semantics | L | AG, mid-to-posterior STS,<br>posterior MTG/ITG | -56 | -38 | 2 |
|  |  |  | -46 | -66 | 26 |
|  |  |  | -48 | -54 | -14 |
|  |  |  | -44 | -44 | -18 |
|  |  |  | -62 | -20 | 2 |
| Theory of mind | L | AG, mid-to-posterior STS | -52 | -58 | 22 |
|  |  |  | -56 | -42 | 4 |
|  |  |  | -58 | -26 | -6 |
|  |  |  | -54 | -52 | 8 |
|  | R | AG | 54 | -54 | 24 |
| Tools | L | Posterior MTG | -50 | -66 | -6 |
|  |  |  | -52 | -58 | 2 |

### Left hemisphere

In the left hemisphere, significant clusters were found across six domains, with only faces failing to show significant activation likelihood in this lateral region (see Figure 1).

The semantics domain engaged a large region of the left hemisphere, extending from the edge of the AG, along the posterior MTG/STG/STS, toward the anterior edge of the ROI (the whole brain results reported by Jackson ^8^ extend into anterior temporal lobe) and ventrally into posterior ITG. These posterior MTG and ITG regions were also found to be activated consistently across the semantic control assessments, demonstrating the particular role of the ventral aspects of this region for the controlled use of semantics, whilst the more dorsal regions identified only in the general semantics contrast may reflect more general semantic processes (consistent with Jackson ^8^). The phonology domain was associated with consistent recruitment of the mid-to-posterior STG, as well as a region of posterior ITG overlapping semantics and semantic control. The theory of mind cluster was similar in extent and volume to the semantics cluster, extending from AG along the STS until it reached the anterior edge of the ROI, although lacking involvement of the ventral portion of MTG and ITG areas associated with control. Note that either inferior parietal or posterior temporal regions could be referred to as TPJ when studying theory of mind ^45^. Smaller clusters were identified for tools and biological motion in the most posterior portion of the ROI near the temporo-occipital border. Biological motion consistently recruits the most posterior region of any domain, located in the pMTG bordering the middle occipital gyrus, just dorsal to the tools cluster. Tools consistently recruit an area of the middle temporal sulcus near the temporo-occipital junction, inferior to biological motion and overlapping semantic control.

Formal ALE contrast analyses were conducted for each pair of domains with overlapping voxels. As the overlapping nature of semantics and semantic control is expected and established ^8^, the contrasts between semantics and semantic control, and semantics and tools (in the semantic control regions shown to overlap with tools below) do not aid the results interpretation further and can be found in the Supplemental Materials (see Supplemental Figures 14-16). Formal conjunctions and contrasts for the remaining pairwise comparisons are displayed in Figure 2.

**Figure 2.**
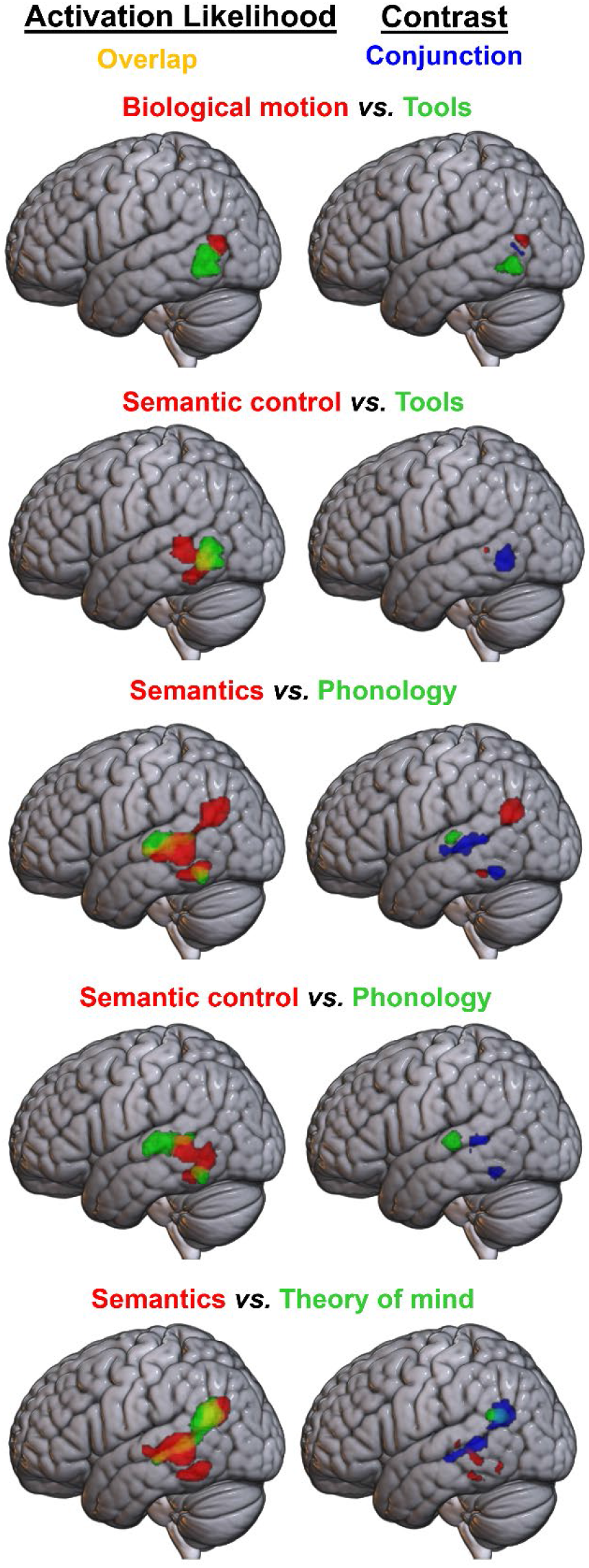
Formal contrast analyses between pairs of domains with overlap in the left hemisphere at a voxel-level threshold of p<.001. Left column: pairwise overlays of ALE maps, showing domain A in red and domain B in green, with overlap in yellow. Right column: results of formal contrast and conjunction analyses; domain A > B is shown in red, B > A in green, and their conjunction in blue.

The contrast analyses reveal the possible relations between tools and the other domains. Although the tools cluster is bordering the biological motion result, direct comparison revealed little conjunction between these domains, indicating distinct regions. However, the majority of the tools result overlaps with the dorsal aspects of the semantic control cluster without differential involvement, and there are no voxels showing significantly greater involvement in tools over semantic control. This could reflect a single region for semantic control, which may be engaged for any semantic content but is engaged to a greater extent for tools than other stimuli (see the Discussion for consideration of why this could be the case).

While the phonology and semantics (and to a lesser extent semantic control) clusters overlap along their edges in the STS, the STG involvement was unique for phonology. The ventral edge of AG and a small region of pITG demonstrated greater involvement for semantics. This suggests relatively distinct regions of pLTC for semantics and phonology, particularly as the semantic domain includes auditory words, so the influence of phonology cannot be entirely excluded. Both phonology and semantic control engaged the pITG. Although the pMTG was identified in semantic control only, this difference did not reach statistical significance.

Direct contrasts revealed a large region of overlap between semantics and theory of mind. Despite this, inferior (semantic control-related) pLTC areas demonstrated greater semantic involvement.

Although the more dorsal parietal region is found consistently for both semantics and theory of mind, it has a greater likelihood of activation within the theory of mind domain.

### Right hemisphere

Significant consistent recruitment of right hemisphere pLTC regions was found in only four domains: faces, theory of mind, biological motion and phonology. Further ALE analyses were conducted for subsets of three of these domains, to minimise the possible confounding effects of overlap in the content of the included studies (see Figure 3 and Table 2).

**Figure 3.**
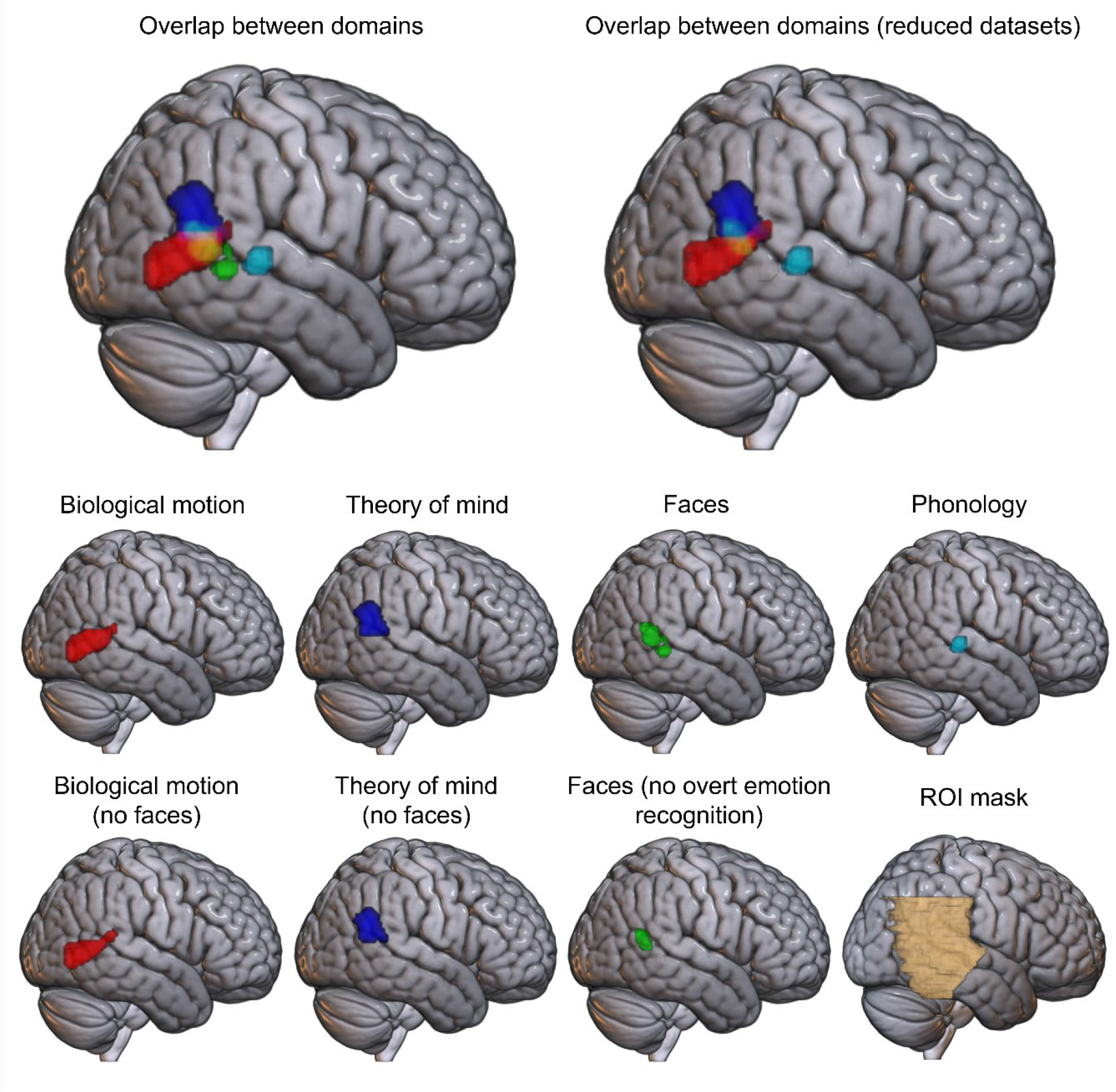
Activation likelihood estimation maps for the four domains with significant results in the right hemisphere. Top row: overlap between domains, with and without exclusions designed to minimise overlap in the included content. Middle row: regions consistently identified across studies in the full dataset for each domain. Bottom row: results of reduced datasets designed to minimise overlap in the content included within each domain. Bottom right: the right hemisphere aspects of the region of interest mask used for all analyses. Domains not shown have no significant results in the right hemisphere.

**Table 2.** Activation likelihood estimation across reduced datasets in the right hemisphere

| Domain | L/R | Region of Activation | Peak MNI Coordinate |  |  |
| --- | --- | --- | --- | --- | --- |
|  |  |  | x | y | z |
| Biological motion | L | Posterior MTG | -50 | -68 | 8 |
|  | R | Posterior MTG | 50 | -68 | 0 |
|  |  |  | 50 | -64 | 2 |
|  |  |  | 60 | -42 | 16 |
| Faces | R | Posterior MTG | 54 | -56 | 16 |
|  |  |  | 54 | -52 | 10 |
| Theory of mind | L | AG | -52 | -58 | 22 |
|  | R | AG | 54 | -54 | 24 |

Phonology engaged a small region of the posterior middle STS, a homologue of the region engaged in the left hemisphere. This region did not demonstrate overlap with any other domain. The remaining three domains all engaged posterior aspects of the pLTC, yet differed on the ventral-dorsal dimension. Theory of mind consistently recruited the AG and posterior STG. Biological motion engaged the posterior aspects of the temporal lobe, including MTG, ITG and to a lesser extent, STG. A small region of posterior MTG/STS was implicated in the faces domain.

Three domains had overlapping voxels in the right hemisphere: theory of mind, biological motion and faces (see Figure 4 and Table 2). As theory of mind engaged a more dorsal region than biological motion, their overlap was minimal at the edge of each cluster, suggesting distinct, albeit nearby areas are implicated in these domains. However, the faces cluster was intermediate between theory of mind and biological motion, showing considerable overlap with both. To assess whether this overlap was the result of the use of face stimuli in some studies within the biological motion and theory of mind domains, reduced datasets excluding face stimuli were considered. Removing face stimuli had little effect; the studies employing face stimuli are insufficient to explain the overlap between theory of mind or biological motion and faces. In addition, the overlap between theory of mind and faces could be hypothesised to be due to the use of emotional recognition tasks in the face domain, a task used to assess both face processing and theory of mind. However, excluding experiments featuring explicit emotion evaluation tasks from the faces domain, did not reduce this overlap. Indeed, although fewer studies resulted in a smaller cluster, it was the regions that overlapped with biological motion and theory of mind that remained. Thus, these potential confounds cannot explain the identified organisation of right pLTC, with substantial overlap between faces and both theory of mind and biological motion, even after controlling for these effects.

**Figure 4.**
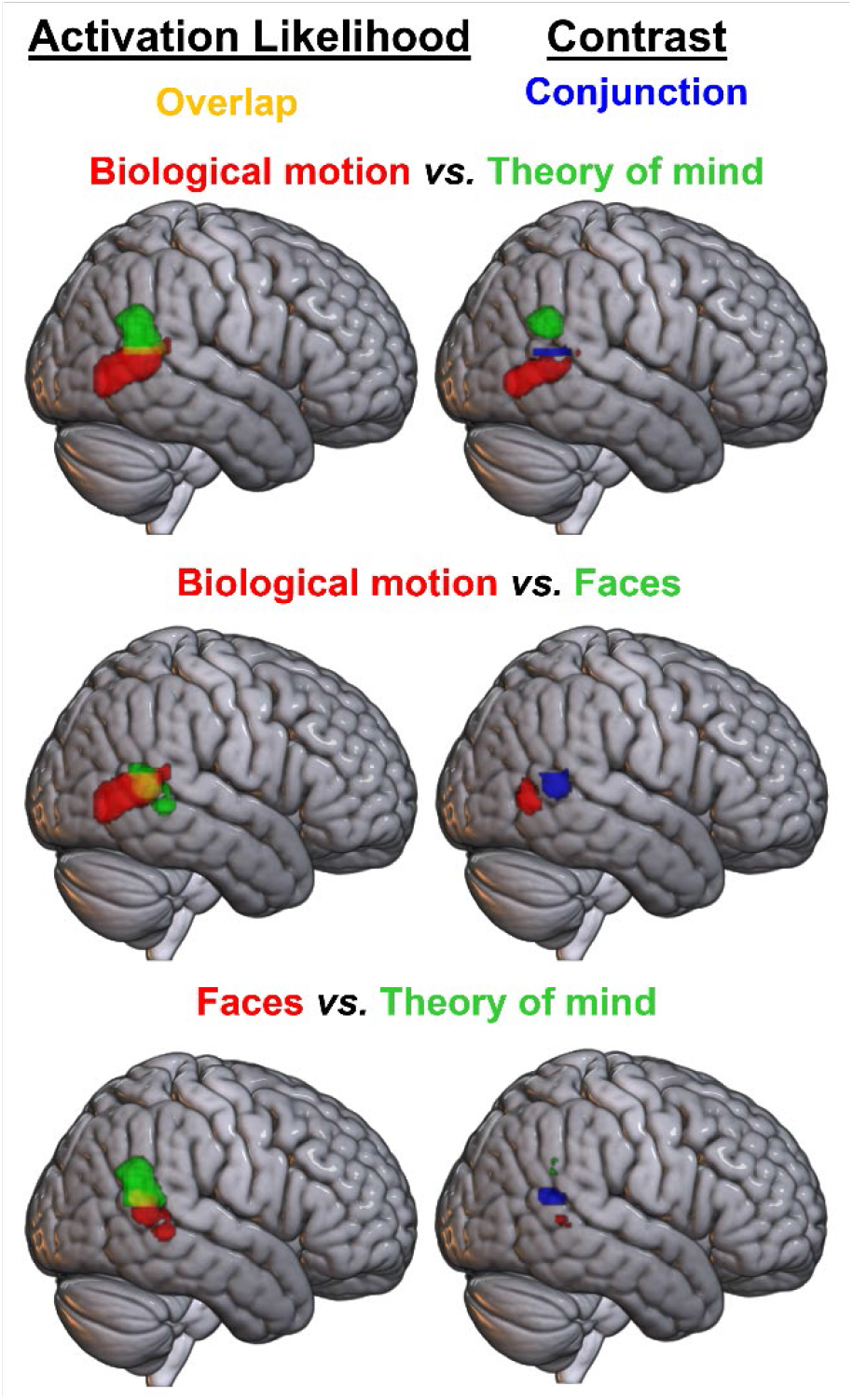
Formal contrast analyses between pairs of domains with overlap in the right hemisphere at a voxel-level threshold of p<.001. Left column: pairwise overlays of ALE maps, showing domain A in red and domain B in green, with overlap in yellow. Right column: results of formal contrast and conjunction analyses; domain A > B is shown in red, B > A in green, and their conjunction in blue.

The results of pairwise formal contrast analyses between these domains are shown in Figure 4 (also see Table 4). As the cross-domain overlap could not be explained by any of the factors assessed in the reduced datasets these formal contrasts employed the full datasets to maximise power. Formal contrasts between reduced datasets gave similar results and can be found in the Supplemental Materials (Supplemental Figures 14-16). Contrast analyses demonstrated significant differences in activation likelihood across the majority of the biological motion and theory of mind clusters suggesting relatively distinct regions for biological motion, in the most posterior part of the MTG at the border of the occipital lobe, and theory of mind, in the AG. Although overlapping with the faces domain, areas in each of these subregions displayed significantly greater activation likelihood for their associated domain than for faces. Whilst a small region of posterior MTG was identified for faces over theory of mind, this fell within the biological motion area and there were no significant results for faces over biological motion. This provides no support in favour of there being face-specific (exclusively involved in faces) pLTC subregions, although some areas may show relatively greater responses to face stimuli.

**Table 3.** Pairwise contrast analyses in the left hemisphere

| Contrast | Region of Activation | Peak MNI Coordinate |  |  |
| --- | --- | --- | --- | --- |
|  |  | x | y | z |
| Biological motion > tools | V5/MT | -48 | -70 | 8 |
|  |  | -48 | -68 | 14 |
|  |  | -50 | -66 | 10 |
| Tools > biological motion | ITG | -54 | -66 | -12 |
|  |  | -48 | -52 | -8 |
| Semantic control > tools | MTG | -54 | -44 | 2 |
| Semantics > phonology | AG/MTG | -46 | -62 | 20 |
|  |  | -42 | -66 | 20 |
|  | Fusiform/ITG | -38 | -16 | 0 |
| Phonology > semantics | MTG | -56 | -24 | 0 |
|  | STG | 58 | -28 | 0 |
|  |  | 60 | -32 | 0 |
| Semantic control > phonology | No peaks |  |  |  |
| Phonology > semantic control | MTG | -64 | -24 | 2 |
| Semantics > theory of mind | ITG | -60 | -46 | -10 |
|  | ITG | -46 | -40 | -20 |
|  | ITG | -48 | -60 | -12 |
|  | MTG | -64 | -30 | 0 |
| Theory of mind > semantics | MTG | -54 | -62 | 20 |

**Table 4.**
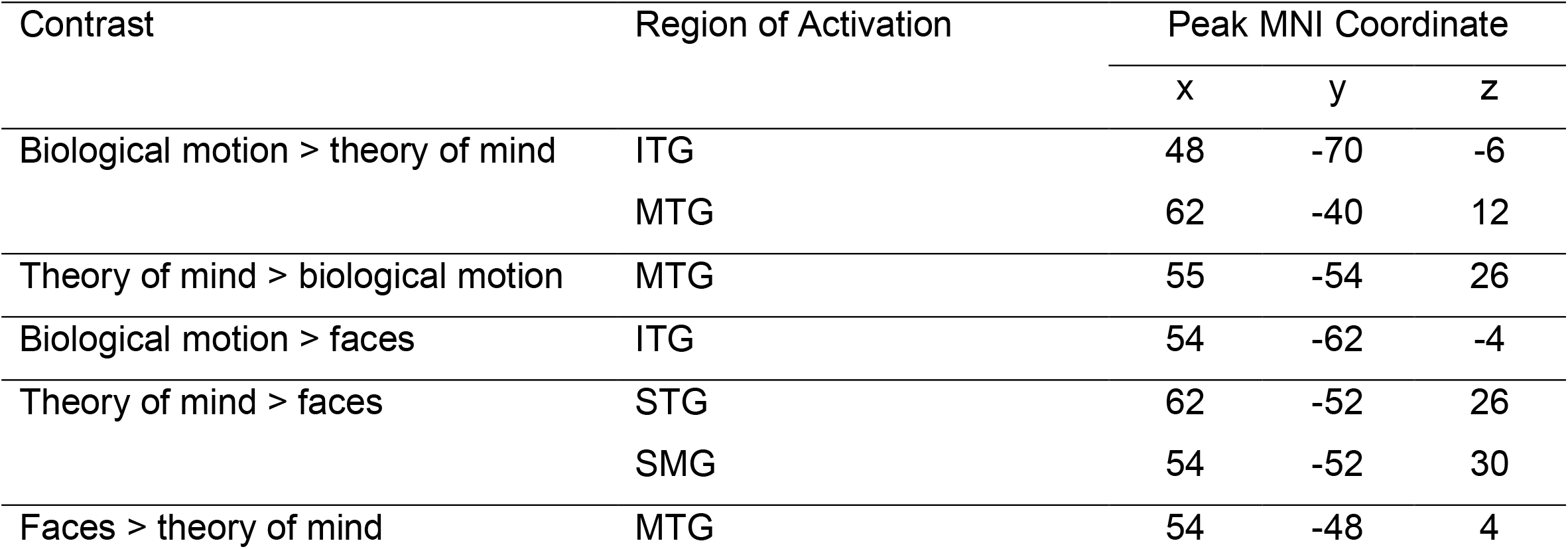
Pairwise contrast analyses in the right hemisphere

## DISCUSSION

Direct comparison across a series of seven ALE meta-analyses assessing diverse domains, delineated the functional organisation of the pLTC. The resulting structure was neither a single discrete region per domain (as previously associated with the basal temporal lobe ^47^) nor a highly overlapping region suggesting a great deal of shared processing across many domains (as identified in the inferior parietal cortex ^56^), but a midpoint on this continuum. Many domains recruited dissociable areas, likely reflecting discrete functional regions. However, a number of domains implicated highly similar subregions and may reflect shared processing. These findings have been synthesised into a summary diagram (Figure 5). The remainder of the Discussion considers this organisation and its implications, including possible explanations for the reliance of multiple domains on shared regions.

**Figure 5:**
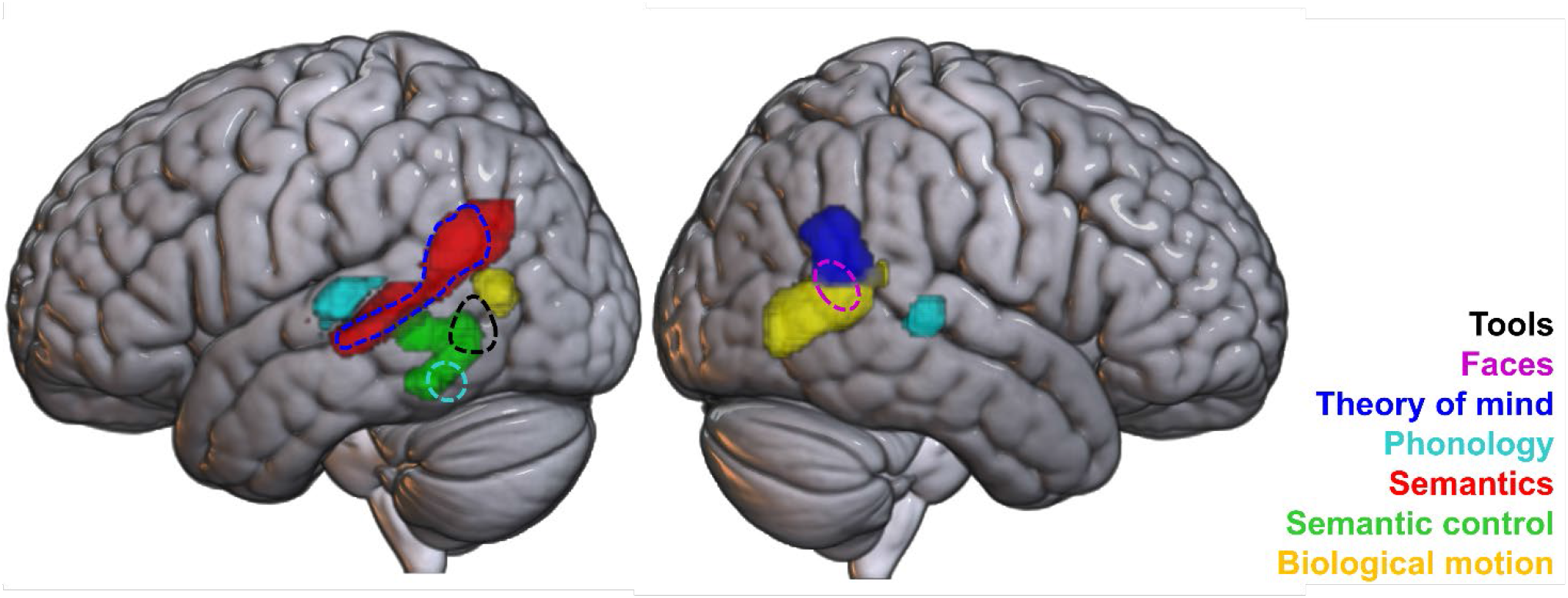
A synthesis of results across the seven domains in the bilateral posterior lateral temporal gyrus ROI. Regions with distinct activation are seen for some domains, e.g., phonology and biological motion, while some domains appear subsumed by others, such as tools by semantic control (indicated by dashed lines).

### Left-lateralised domains

#### Semantics and phonology

Both semantics and phonology had a left hemisphere focus (albeit with some additional right pLTC involvement for phonology only). These domains were underpinned by dissociable pLTC regions consistent with previous meta-analytic, theoretical and neuropsychological work ^7,8,11,32–34,68–72^. A discrete region of bilateral STG was implicated in phonology; this region is associated with processing speech sounds ^73,74^ and damage here is linked to conduction aphasia ^36–38^. In contrast, a swathe of the left pSTS extending dorsally to the ventral edge of AG, was implicated in general semantic processing (due to its involvement in semantics, yet not semantic control) which overlapped only minimally with phonology. This indicates broadly separable regions for phonology and semantic representation in the pLTC, consistent with the relative independence of damage to semantic and phonological processes observed within the neuropsychological literature ^75,76^. For instance, while damage to the pSTG leads to the combined semantic-phonological impairment seen in Wernicke’s aphasia, lesions in the pMTG result in multimodal semantic impairment without phonological impairment, seen in semantic aphasia ^75^. This dissociation is consistent with sharper cytoarchitectural distinctions between the STG and MTG than between other temporal gyri ^77–79^.

The precise interpretation of the semantic pSTS region, which is seen in the present analysis for semantics but not phonology or semantic control, requires further investigation. One possibility is that the pSTS region may be involved in the comprehension of sentences ^80,81^ or of narrative gestalt meaning; this region is more activated as consistent, time-extending meaning builds up ^82^.

#### Semantic control

The left pITG/pMTG was also associated with semantics, specifically the controlled access and manipulation of semantics. This aligns with neuroimaging studies of semantic control ^2,7,8,55,83^, and with the regions typically damaged in semantic aphasia, in which patients have difficulty accessing weaker or subordinate associations, and poor inhibition of strong associations, following damage to this area due to stroke ^3,5,84^. A posterior lateral temporal region for semantic control is also consistent with modelling work that indicates a need for control processes to interact with modality-specific spokes, rather than the multimodal ATL hub ^85^. The pMTG/pITG may act as an intermediary between the ATL hub and the IFG ^86^, and is well-placed to interact with surrounding modality-specific spokes, such as visual areas in the fusiform gyrus and occipital lobe, auditory areas in the STG, and praxis areas in inferior parietal cortex ^1,8,69^.

An overlapping region of left pITG was also implicated in phonology. Previously demonstrated to respond particularly to hard phonological tasks ^55^ and implicated in the extended multiple demand network ^87^, this region may reflect shared control processing for language subdomains or across a broad set of domains. It is unclear whether this constitutes a discrete region, or whether pMTG and pITG demonstrate a graded shift in a preference for the control of meaningful stimuli. The distinction between control and representation processes seems to be a key organisational principle for language regions ^8,55^ and the brain more broadly ^88–90^, and as such is a critical factor to consider when delineating the organisation of the pLTC. Representation regions engage more dorsal subregions of the left posterior temporal lobe, while control processes – which may be domain-general, or specific to language – engage more ventral subregions, such as the pMTG and pITG ^8,55,87^.

#### Tools processing

The tools domain also overlaps the semantic control cluster, in a pMTG region more dorsal and posterior to, and thus distinct from, the pITG region identified for phonology. Tools are one semantic category amongst many, and the present analyses found no evidence that tools recruit a distinct region beyond those that are implicated in semantics across categories; however, it is notable that tools engage an area specialised for semantic control, rather than semantic representation. Whilst differences in the relative engagement of basal temporal areas across categories are hypothesised to relate to low level visual properties of the stimuli ^91–95^, differences in the lateral temporal cortex may have alternative explanations related to higher-level multimodal processes. In comparison to other semantic categories, tools may rely more heavily on control processes ^10^. Much of our conceptual understanding of a manipulable tool relies on praxis, which requires the unfolding of a sequence of movements across time and space. This dynamic time-varying information may be more complex than the static feature information that is sufficient for comprehension of most other semantic categories, such as size or shape, and as such may require greater control to selectively access and manipulate the relevant features. This region may perform semantic control for all categories with tools simply requiring a high level of semantic control, or it may have a particular role in integrating information between the praxis network for planning tool use (in the superior parietal lobule and premotor cortices) and the more conceptual temporal lobe system for tool knowledge ^13,96^.

#### The relationship between theory of mind and semantics

Theory of mind recruited the same left pLTC subregions as those recruited for semantic representation, including the pSTS and ventral AG. This area is often labelled as the TPJ in theory of mind literature, a region considered to be of particular importance for mentalising ^42,44,57,97,98^. It has previously been established that the theory of mind network includes semantic regions ^99–103^; here, by formally comparing across domains, we have demonstrated that the same regions are recruited for both domains, a similarity that may typically be obscured by the inconsistent use of anatomical terms or the tendency to investigate each domain independently. There are many possibilities for this high degree of overlap in the left pLTC. Many theory of mind tasks involve meaningful stimuli, such as narratives or vignettes, which would necessarily engage semantic processing areas to comprehend the meanings of the words and track meaning – or indeed multiple meanings – over time ^57,82^, and so this overlap may simply reflect a need for the engagement of semantic networks in theory of mind processing ^102,103^. These activations may have been detected in the present meta-analysis due to different theory of mind task conditions not being adequately matched on semantic demands.

Direct contrasts reveal that whilst the ventral anterior AG is engaged for both domains, activation likelihood is significantly higher for theory of mind than semantics in this region. Furthermore, the right AG was identified for theory of mind but not semantic cognition. This may suggest that the AG is responsible for theory of mind and some of the semantic studies engage relevant processes. However, the majority of semantic studies that were included utilised single words, which should not require theory of mind processing. Indeed, the role of the AG is a focus of great debate within the semantic cognition literature ^7,104–107^ and the laterality of the semantic network is task- and stimuli-dependent ^108,109^. Similarly, whilst theory of mind is often considered to rely on right-lateralised areas ^110,111^, it may be that these laterality differences are present only after subtracting a left-lateralised verbal semantics network through the use of a non-mentalising story in false belief tasks. Thus, a more nuanced explanation may require consideration of multiple different factors affecting both the recruitment of the AG and the laterality of the network engaged, such as the use of sentences ^82,112^, the stimulus modality and the effect of difficulty-dependent deactivation ^113–115^, as well as the social nature of the stimuli. Indeed, as the theory of mind dataset contains a large number of tasks with full sentences, while the semantics dataset contains many tasks that focus on single words, this region may show relatively greater activation for theory of mind on the basis of stimuli differences and not theory of mind requirements. If this is viewed as one large bilateral network performing both kinds of tasks with variations in the locus of peak activation, a very different picture emerges of the regions responsible for these domains. The overlap between these two domains demonstrates the clear need for cross-domain comparisons and careful consideration of terminology. It may be that the questions surrounding the AG and the pLTC in the theory of mind and semantic literature form two parts of a single puzzle where progress would be best achieved by bringing these domains together in future research.

### Right-lateralised domains

Other domains were more likely to engage the right pLTC. Biological motion had a right hemisphere focus engaging a distinct region ventral to theory of mind, centred on the most posterior aspect of MTG (posterior to the other left hemisphere pMTG subregions), immediately ventral to the AG and anterior to the occipital lobe. This placement is consistent with prior assessments identifying a region in pSTS for processing dynamic biological stimuli, and forms part of the dorsal route for visual input, with biological motion processing necessitating connections from occipital cortex, including the extrastriate body area (a region demonstrating preferential responses to static body stimuli) ^19–21,23,116–124^. It is also notable that the overall pattern of lateralisation in the biological motion and faces domains is consistent with the suggestion that social domains are typically right-lateralised ^41,110,125–127^, and with connectivity analyses that find the right pSTS to be more strongly connected to other biological motion and face processing regions in the right hemisphere, than the left pSTS is to its counterparts ^125^.

Although distinct regions, the proximity of theory of mind and biological motion areas may support crucial interactions between these domains. Both of these regions demonstrate partial overlap with a small region identified for faces around their border in the right pSTS. This corresponds to a known face-responsive area that is specialised for processing changeable aspects of a face, such as expression, lip movement and eye gaze ^16–18,26,27^, although this region is typically reported to be bilateral ^16,17,61^. The right-lateralisation in the present analysis could be due to biases in the literature, such as focusing on the right hemisphere for faces or assessing non-speech facial expressions and movements ^128^. This face region falls entirely within areas implicated in theory of mind and biological motion without displaying significantly greater engagement for faces. Indeed, when revisiting the prior literature with this result in mind, it is not clear that this face pSTS region has been reliably differentiated from the proposed pSTS region for biological motion ^23,129,130^. It is possible that there are distinct neural populations subserving each domain, but they cannot be disentangled with meta-analyses or group-level analyses. Alternatively, the engagement of this region for faces may simply reflect the common engagement of biological motion and theory of mind processes in studies employing face stimuli. The processing of dynamic and changeable aspects of faces may be subserved by the pSTS as a subset of all dynamic body stimuli. Thus, this region may not be specialised for faces exclusively, but instead for the processing of dynamic biological stimuli in ventral aspects and for information about intentional action more broadly in dorsal aspects, of which expression and facial movements are a subset. The extensive connectivity of the pSTS with regions that subserve social processing tasks, such as the perception of salient social stimuli, the observation and understanding of intentional action, and the attribution of mental states could support this interpretation and may suggest a particular role for this region as a hub or interface between networks supporting distinct tasks ^125,131^.

## Conclusions

The application of a cross-domain approach here has helped elucidate the wider organisation of the pLTC, illuminating possible subregions and highlighting previously obscured relationships between different domains. Direct comparison utilising a large amount of data across domains, using the activation likelihood estimation method, allowed for a single high-powered study in contrast to a single task-based neuroimaging study, which would have far smaller samples, may be underpowered ^132,133^, and must rely upon single, often idiosyncratic, tasks. Here, we have been able to include thousands of participants, and capture common activations across the different variations of tasks used to assess a process or domain. Further work should focus on testing the hypotheses generated here within individual participants ^134,135^, to determine to what extent apparent overlapping activation is indeed due to shared regions or processes.

## Author Contributions

All authors conceived the study. VJH and RLJ collected and analysed data, VJH wrote the initial draft and RLJ and MALR revised the manuscript.

## Data and Code Availability

The toolboxes used to analyse the data are freely available from www.brainmap.org/ale. The code used can be found in Supplementary Materials 1; foci files and results files can be found on GitHub https://github.com/Vicki-H/Cross-domain-pLTC.

## Ethics

No new data were collected for this study. All data were taken from published works that reported on samples of consenting participants.

