## Supplemental Information for "The cross-domain functional organisation of posterior lateral temporal cortex: Insights from ALE meta-analyses of seven cognitive domains spanning 9515 participants"

**Supplementary Materials**

**Supplementary Materials 1**

Code used for activation likelihood estimation analysis:

java -cp /C:/gingerale/v3.0.2/GingerALE.jar org.brainmap.meta.getALE2 /C:/domain.txt -mask=/C:/mask.nii -p=.001 -perm=10000 -clust=.001 -nonadd

Code used for contrast analysis:

java -cp /imaging/local/software/gingerale/v3.0.2/GingerALE.jar org.brainmap.meta.getALE2Contrast /C:/domain1_p001_C001_10k_ALE /C:/domain2_p001_C001_10k_ALE /C:/domain1anddomain2_p001_C001_10k_ALE -mask=/C:/mask.nii
-out1=domain1 -out2=domain2 -p=.001 -perm=10000 -minVol=20

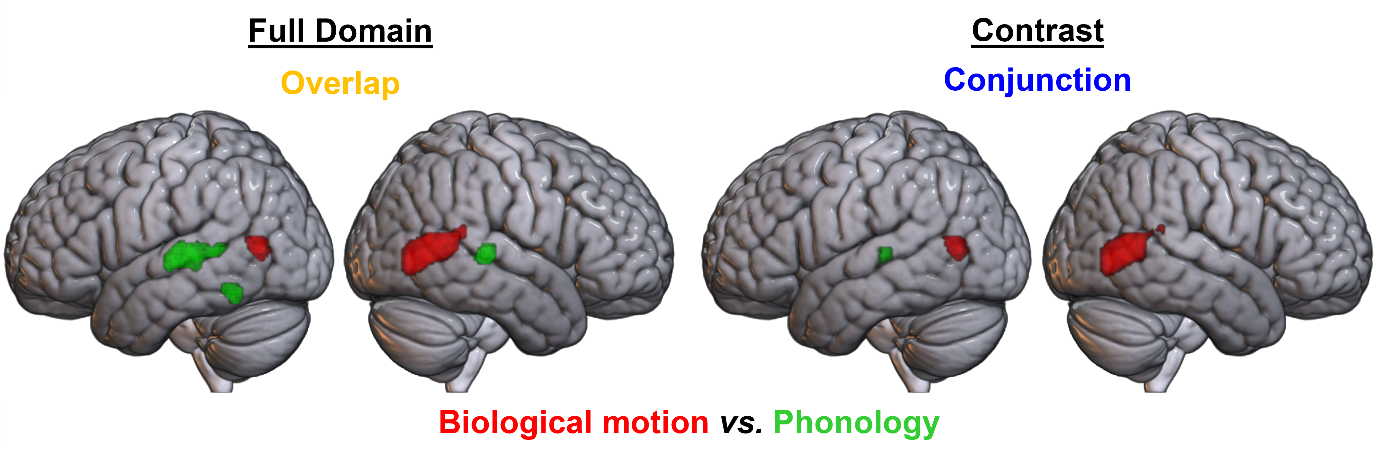

**Supplementary Figure 1.** Formal contrast analyses between the biological motion and phonology domains, at a voxel-level threshold of p<.001. Left: overlays of ALE maps, showing biological motion in red and phonology in green, with overlap in yellow. Right: results of formal contrast and conjunction analyses; biological motion > phonology in red, phonology > biological motion in green, and conjunction in blue.

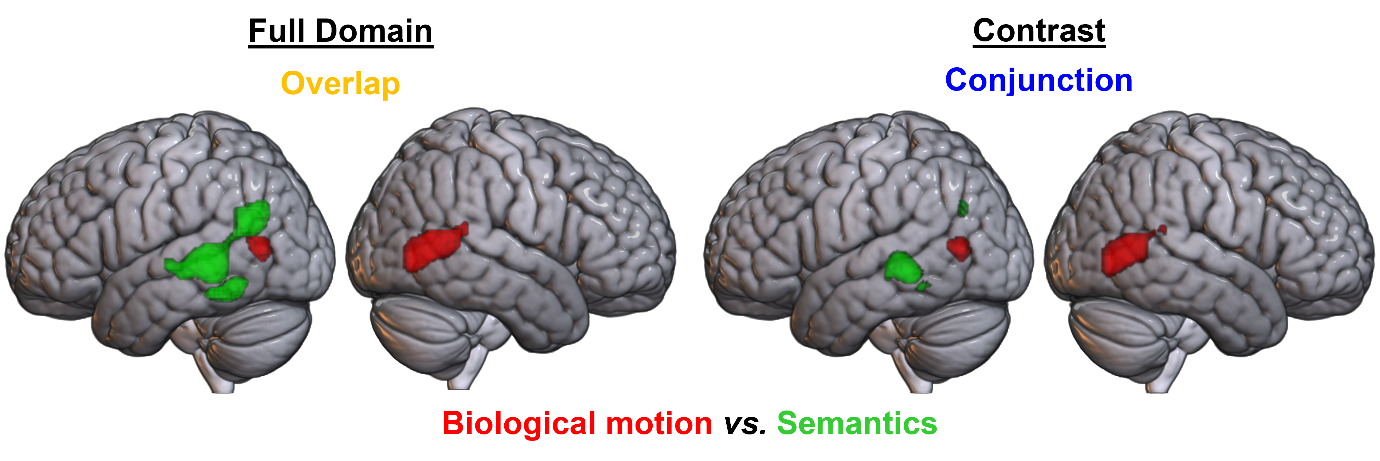

**Supplementary Figure 2.** Formal contrast analyses between the biological motion and semantics domains, at a voxel-level threshold of p<.001. Left: overlays of ALE maps, showing biological motion in red and semantics in green, with overlap in yellow. Right: results of formal contrast and conjunction analyses; biological motion > semantics in red, semantics > biological motion in green, and conjunction in blue.

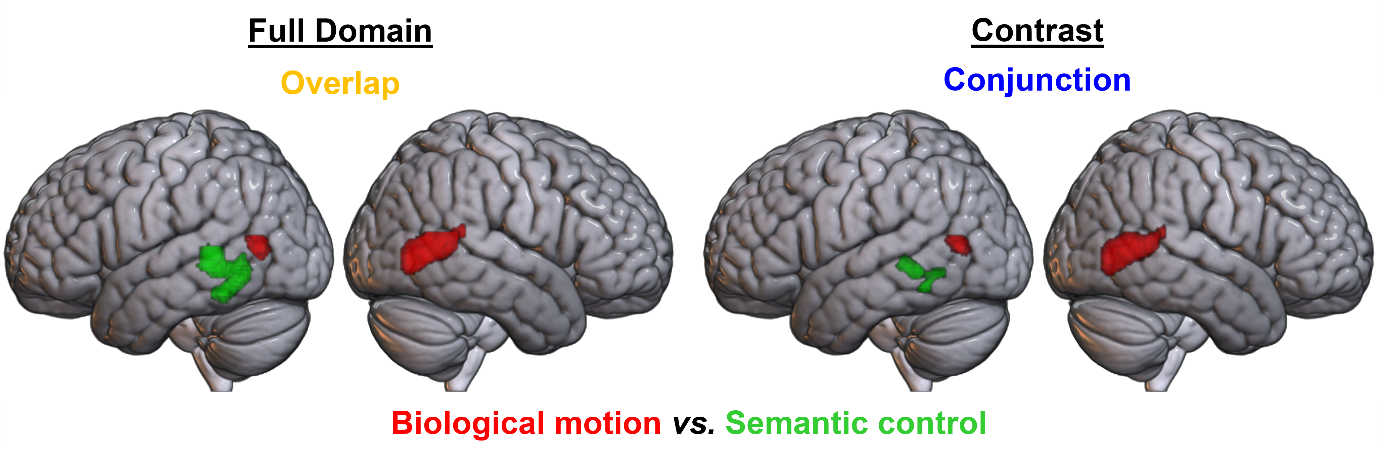

**Supplementary Figure 3.** Formal contrast analyses between the biological motion and semantic control domains, at a voxel-level threshold of p<.001. Left: overlays of ALE maps, showing biological motion in red and semantic control in green, with overlap in yellow. Right: results of formal contrast and conjunction analyses; biological motion > semantic control in red, semantic control > biological motion in green, and conjunction in blue.

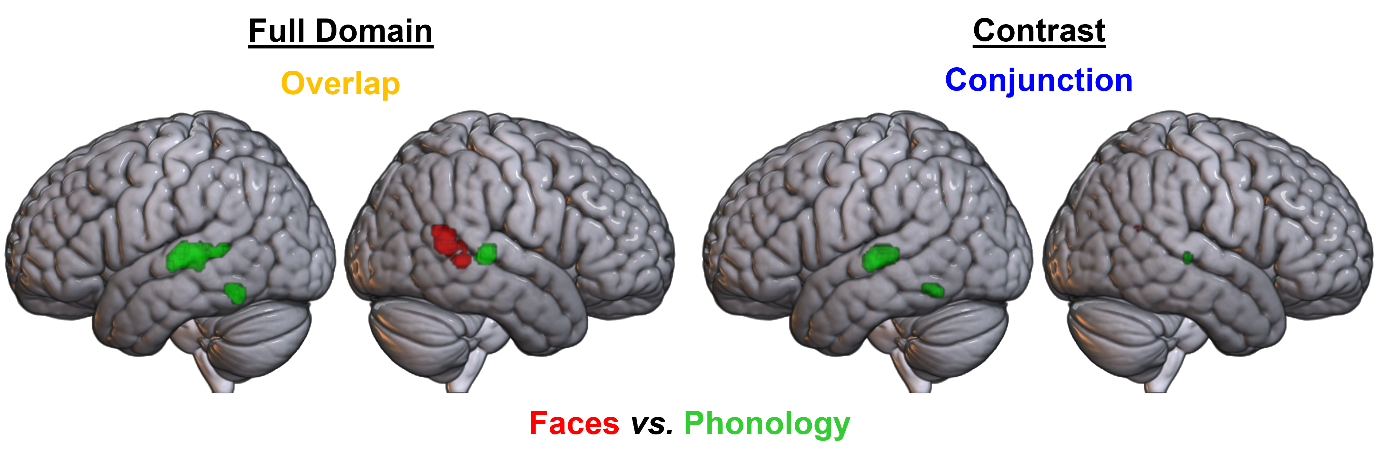

**Supplementary Figure 4.** Formal contrast analyses between the faces and phonology domains, at a voxel-level threshold of p<.001. Left: overlays of ALE maps, showing faces in red and phonology in green, with overlap in yellow. Right: results of formal contrast and conjunction analyses; faces > phonology in red, phonology > faces in green, and conjunction in blue.

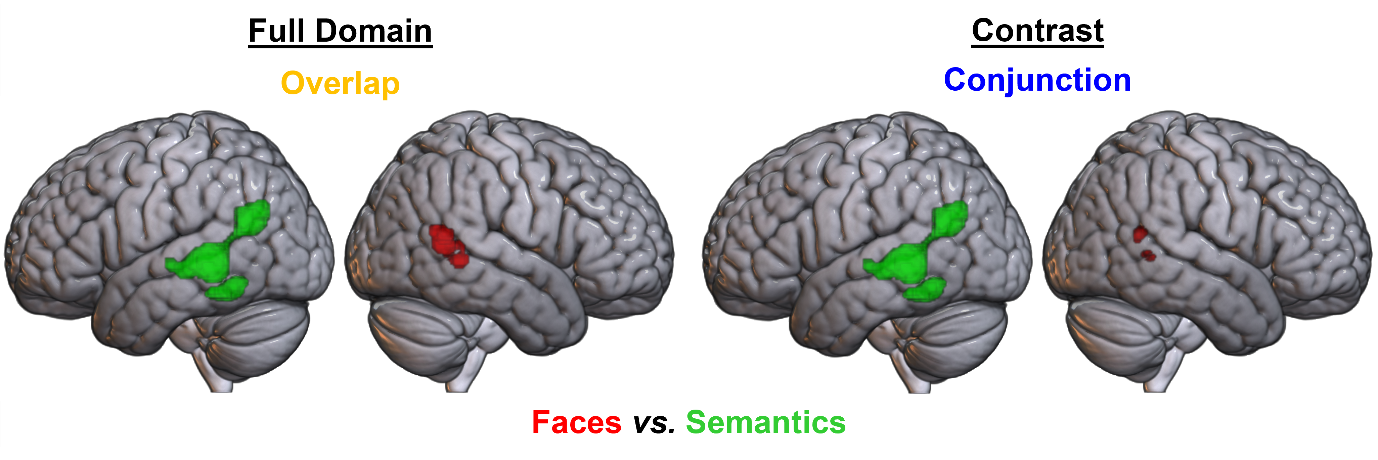

**Supplementary Figure 5.** Formal contrast analyses between the faces and semantics domains, at a voxel-level threshold of p<.001. Left: overlays of ALE maps, showing faces in red and semantics in green, with overlap in yellow. Right: results of formal contrast and conjunction analyses; faces > semantics in red, semantics > faces in green, and conjunction in blue.

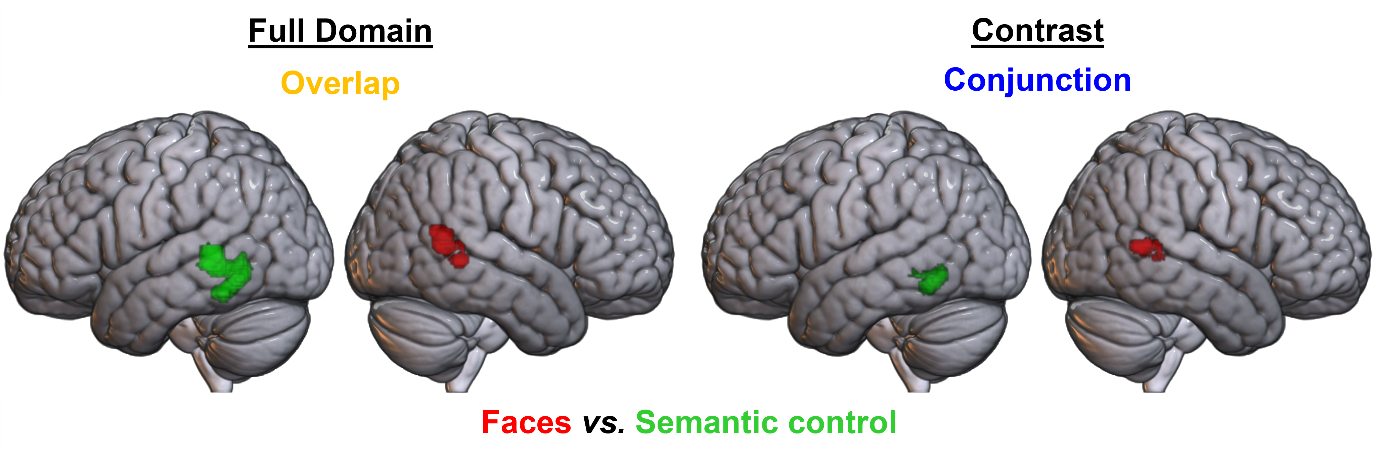

**Supplementary Figure 6.** Formal contrast analyses between the faces and semantic control domains, at a voxel-level threshold of p<.001. Left: overlays of ALE maps, showing faces in red and semantic control in green, with overlap in yellow. Right: results of formal contrast and conjunction analyses; faces > semantic control in red, semantic control > faces in green, and conjunction in blue.

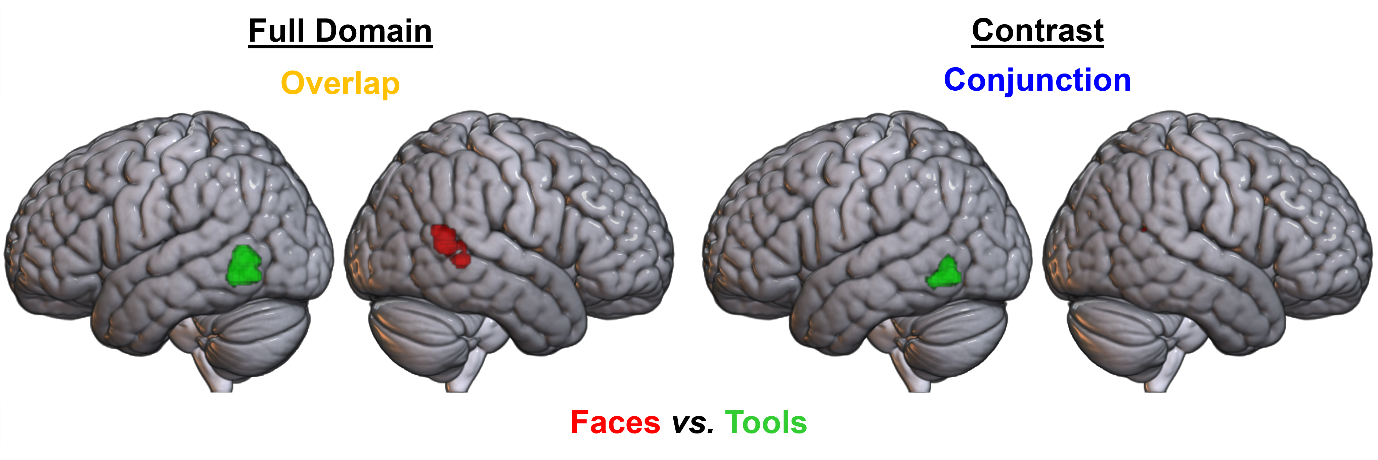

**Supplementary Figure 7.** Formal contrast analyses between the faces and tools domains, at a voxel-level threshold of p<.001. Left: overlays of ALE maps, showing faces in red and tools in green, with overlap in yellow. Right: results of formal contrast and conjunction analyses; faces > tools in red, tools > faces in green, and conjunction in blue.

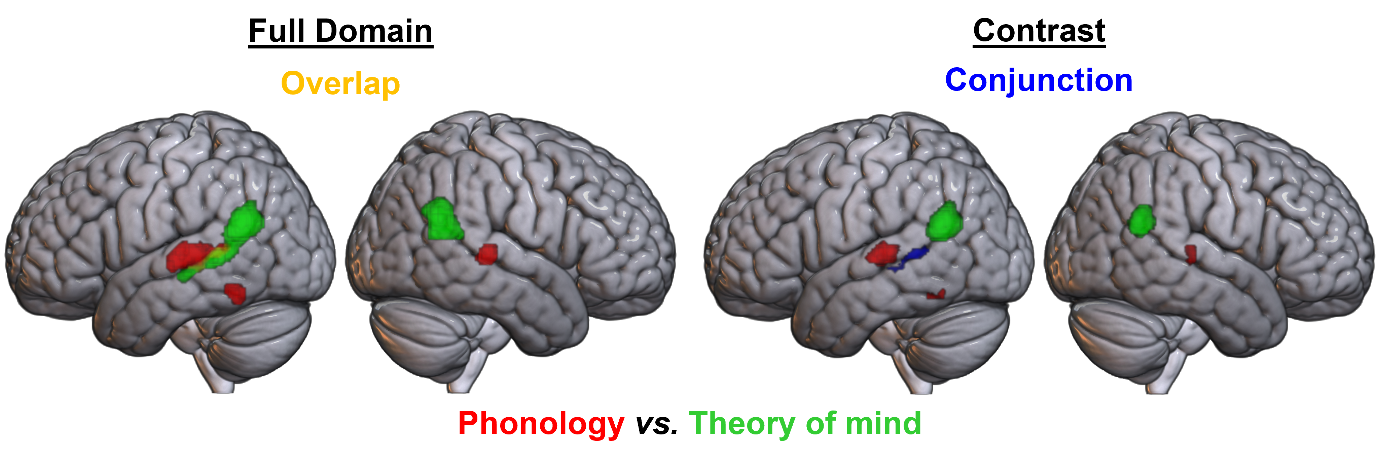

**Supplementary Figure 8.** Formal contrast analyses between the phonology and theory of mind domains, at a voxel-level threshold of p<.001. Left: overlays of ALE maps, showing phonology in red and theory of mind in green, with overlap in yellow. Right: results of formal contrast and conjunction analyses; phonology > theory of mind in red, theory of mind > phonology in green, and conjunction in blue.

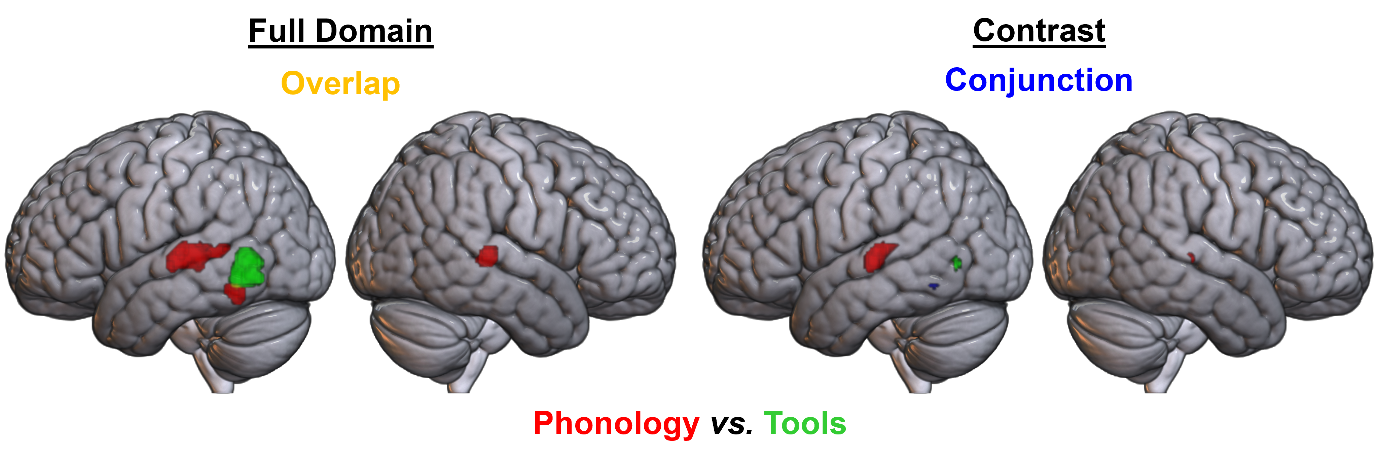

**Supplementary Figure 9.** Formal contrast analyses between the phonology and tools domains, at a voxel-level threshold of p<.001. Left: overlays of ALE maps, showing phonology in red and tools in green, with overlap in yellow. Right: results of formal contrast and conjunction analyses; phonology > tools in red, tools > phonology in green, and conjunction in blue.

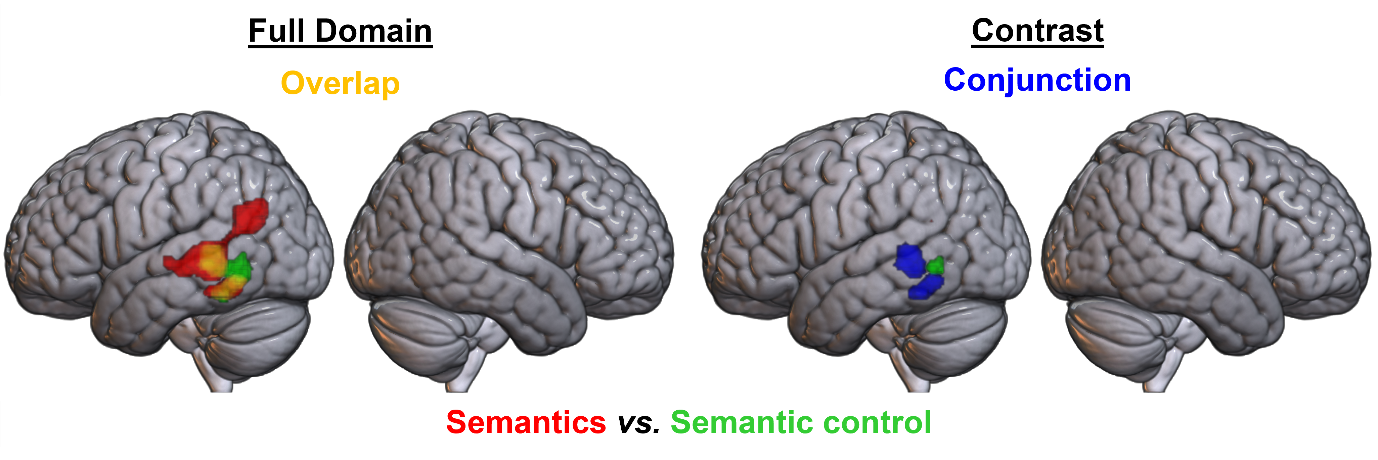

**Supplementary Figure 10.** Formal contrast analyses between the semantics and semantic control domains, at a voxel-level threshold of p<.001. Left: overlays of ALE maps, showing semantics in red and semantic control in green, with overlap in yellow. Right: results of formal contrast and conjunction analyses; semantics > semantic control in red, semantic control > semantics in green, and conjunction in blue.

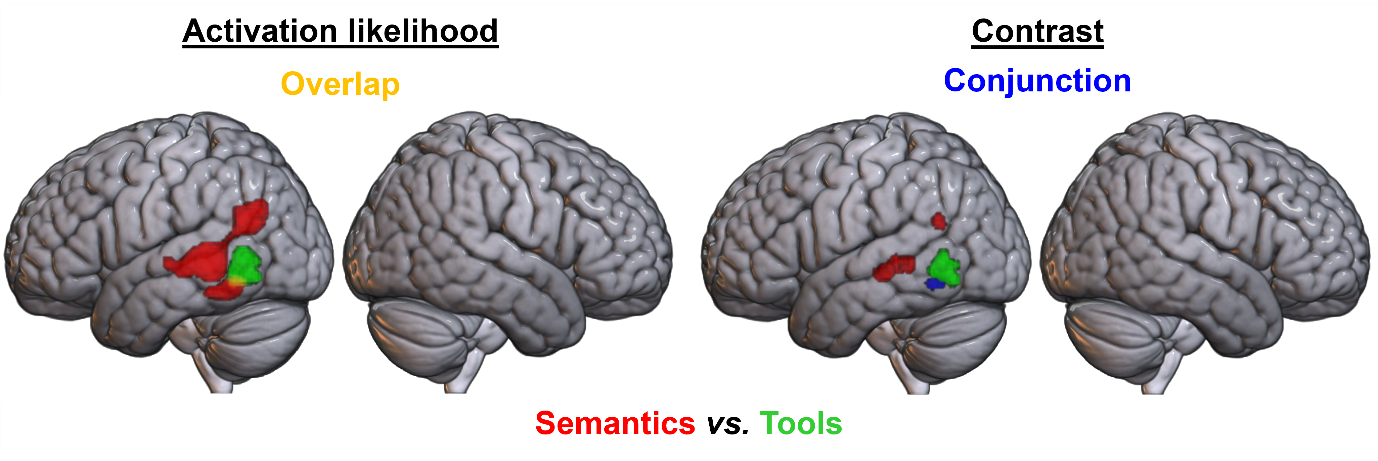

**Supplementary Figure 11.** Formal contrast analyses between the semantics and tools domains, at a voxel-level threshold of p<.001. Left: overlays of ALE maps, showing semantics in red and tools in green, with overlap in yellow. Right: results of formal contrast and conjunction analyses; semantics > tools in red, tools > semantics in green, and conjunction in blue.

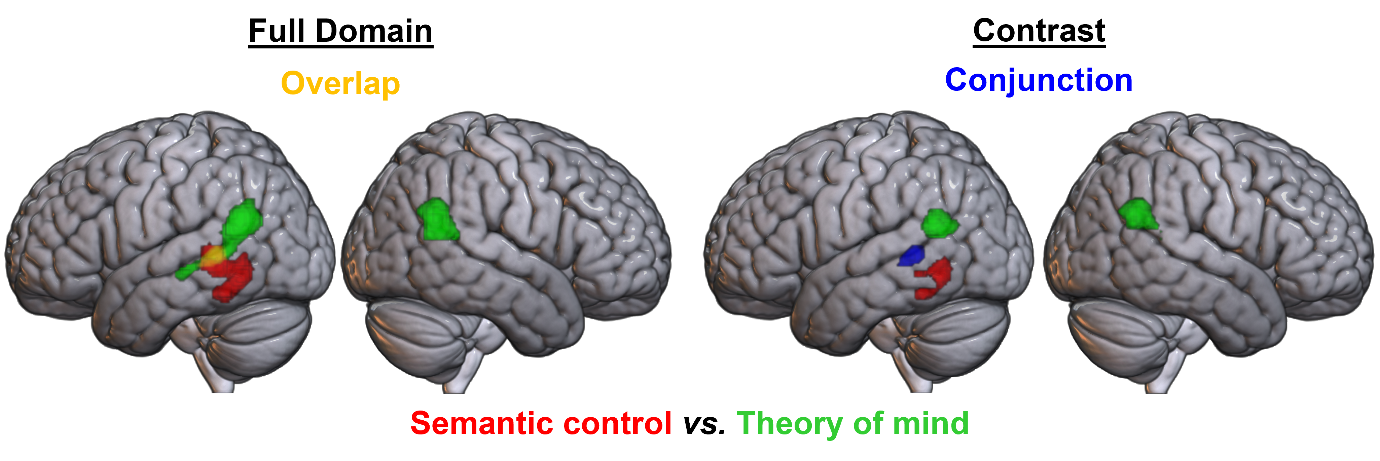

**Supplementary Figure 12.** Formal contrast analyses between the semantic control and theory of mind domains, at a voxel-level threshold of p<.001. Left: overlays of ALE maps, showing semantic control in red and theory of mind in green, with overlap in yellow. Right: results of formal contrast and conjunction analyses; semantic control > theory of mind in red, theory of mind > semantic control in green, and conjunction in blue.

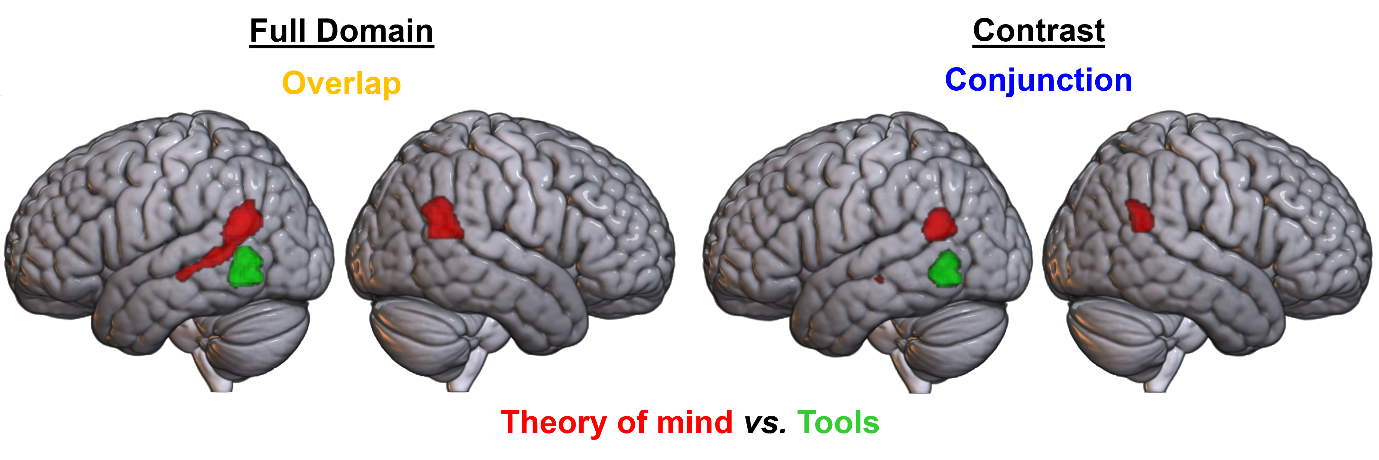

**Supplementary Figure 13.** Formal contrast analyses between the theory of mind and tools domains, at a voxel-level threshold of p<.001. Left: overlays of ALE maps, showing theory of mind in red and tools in green, with overlap in yellow. Right: results of formal contrast and conjunction analyses; theory of mind > tools in red, tools > theory of mind in green, and conjunction in blue.

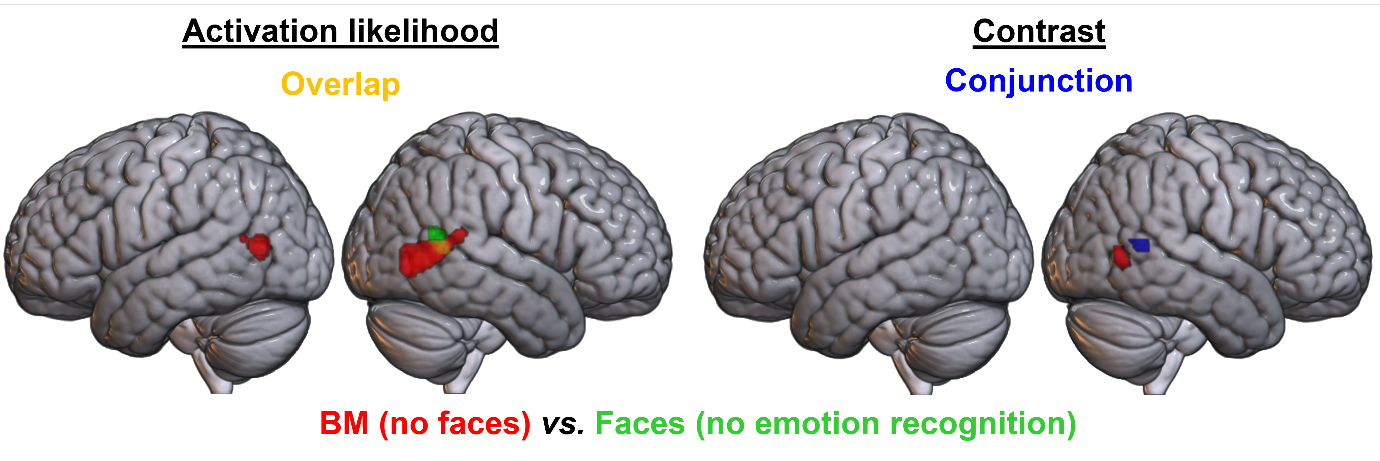

**Supplementary Figure 14.** Formal contrast analyses between the reduced biological motion and reduced faces domains, at a voxel-level threshold of p<.001. Left: overlays of ALE maps, showing biological motion (no faces) in red and faces (no emotion recognition tasks) in green, with overlap in yellow. Right: results of formal contrast and conjunction analyses; biological motion (no faces) > faces (no emotion recognition) in red, faces (no emotion recognition) > biological motion (no faces) in green, and conjunction in blue.

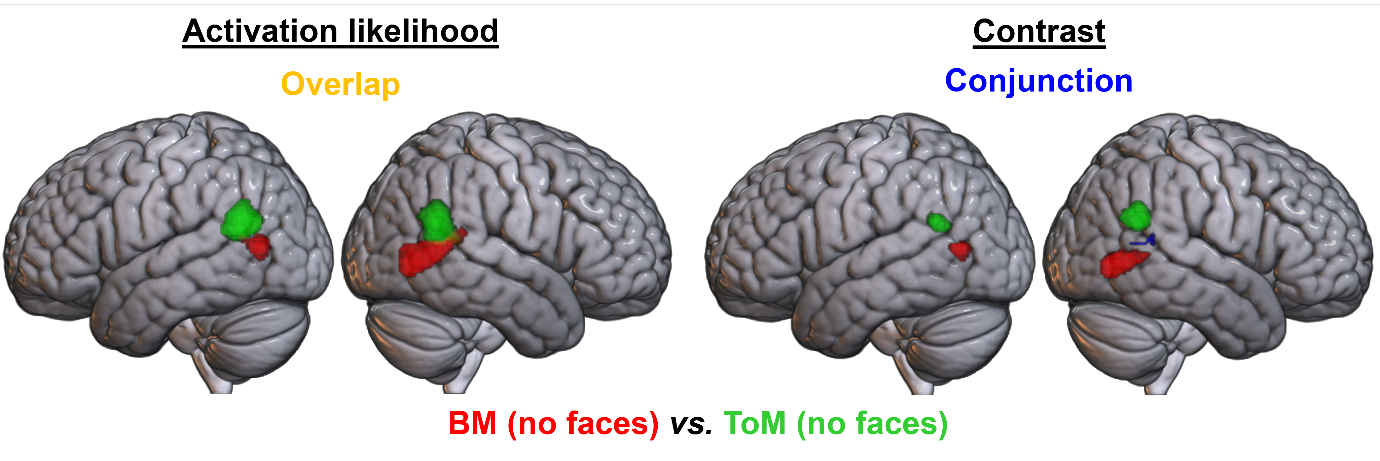

**Supplementary Figure 15.** Formal contrast analyses between the reduced biological motion and reduced theory of mind domains, at a voxel-level threshold of p<.001. Left: overlays of ALE maps, showing biological motion (no faces) in red and theory of mind (no faces) in green, with overlap in yellow. Right: results of formal contrast and conjunction analyses; biological motion (no faces) > theory of mind (no faces) in red, theory of mind (no faces) > biological motion (no faces) in green, and conjunction in blue.

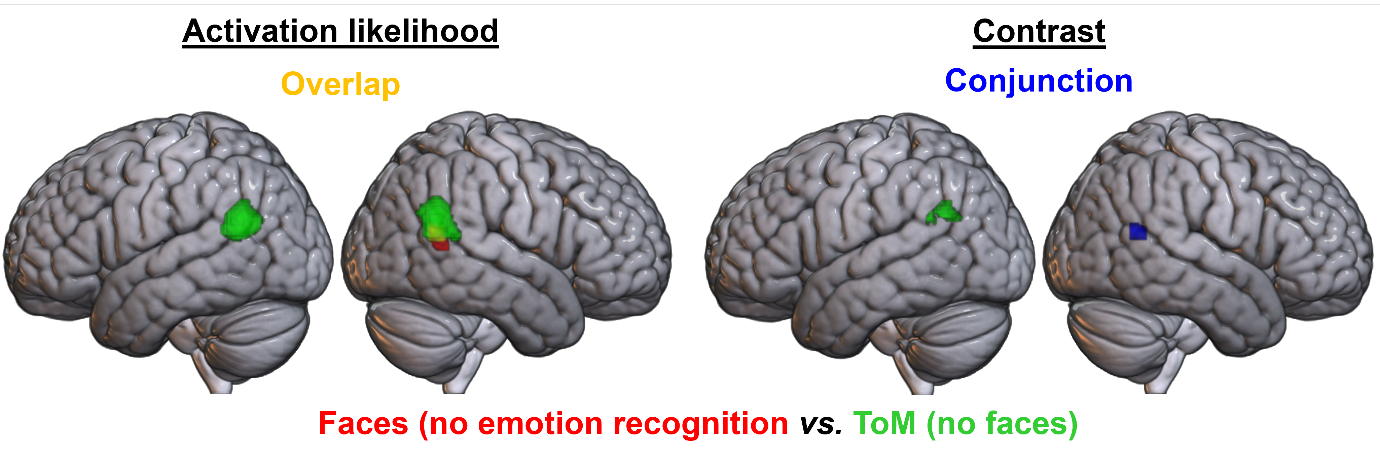

**Supplementary Figure 16.** Formal contrast analyses between the reduced faces and reduced theory of mind domains, at a voxel-level threshold of p<.001. Left: overlays of ALE maps, showing faces (no emotion recognition tasks) in red and theory of mind (no faces) in green, with overlap in yellow. Right: results of formal contrast and conjunction analyses; faces (no emotion recognition) > theory of mind (no faces) in red, theory of mind (no faces) > faces (no emotion recognition) in green, and conjunction in blue.
